# Investigating Rat-Brain Normal Tissue and Tumor FLASH Effects with a Novel Very High Energy Electron Beam

**DOI:** 10.1101/2025.09.05.674499

**Authors:** Tyler V. Kay, Anna L. Price, Markus Sprenger, Victoria J. P. Radosova, Andrew Thompson, Eric L. Martin, Denise Dunn, Victor Popov, Stepan Mikhailov, Zachary J. Reitman, Ying K. Wu, Scott R. Floyd, Mark Oldham

## Abstract

**Purpose:** Ultra-high dose rate (FLASH) irradiation is reported to reduce normal tissue toxicity while maintaining tumor control, however mechanism(s) remain obscure. To study FLASH mechanisms in brain tissue, we developed a novel experimental platform featuring a unique high-energy electron linear accelerator (High Intensity Gamma Source, HIGS) paired with an organotypic *ex vivo* brain metastasis model.

**Methods:** We varied inter-pulse spacing to modulate the mean dose rate (MDR) of our unique 35 MeV electron beam, while maintaining extremely high instantaneous dose rate (IDR), and used film dosimetry to characterize dosimetry and targeting accuracy. We combined this HIGS-FLASH beam with an organotypic rat brain slice/breast carcinoma co-culture model of brain metastasis to assess effects on normal and neoplastic tissues. Live cell and bioluminescence imaging demonstrated cancer cell growth effects, while normal tissue responses and immune activation were assessed with live cell imaging, cytokine profiles, and confocal microscopy. We performed comparison experiments with 20 MeV electrons from a Varian clinical linear accelerator (VCLA) operating at conventional dose rate.

**Results:** The highest IDR of the HIGS-FLASH beam to-date was 20.7 ± 0.6 MGy/s, with maximum MDR of 20.7 MGy/s (1 μs pulse of 20.7 Gy). Beam targeting was accurate to < 1 mm and reproducible. HIGS-FLASH and VCLA dose rates equivalently decreased cancer cell growth. HIGS-FLASH irradiation significantly increased TNFα and fractalkine levels and confocal microscopy revealed distinct changes in microglial morphology in normal brain slices, suggesting microglia activation following HIGS-FLASH irradiation.

**Conclusions:** Our novel experimental platform produces extremely high dose rates and rapid normal/neoplastic tissue readouts for mechanistic research into the effects of FLASH radiation on the brain. HIGS-FLASH irradiation induces comparable cancer cell growth inhibition but differential effects on cytokines and microglial morphology, suggesting that acute innate immune responses may be involved in FLASH normal tissue effects in the brain.

## Introduction

Ultra-high dose rate radiation therapy (FLASH-RT) generally refers to dose rates in excess of 40 Gy/s, which is considerably higher than conventional clinical dose rates (generally 0.1-0.4 Gy/s). Dewey and Boag were the first to report how FLASH-RT affects cell survival. They observed that bacteria were less sensitive to radiation damage with a single high-dose pulse compared with conventional dose rates [1]. Favaudon et al. expanded on this discovery, observing that FLASH-RT reduced fibrosis and apoptosis in mice lungs compared with conventional radiation therapy while simultaneously providing comparable tumor control. This observed effect was named the ‘FLASH effect’ [2]. Reduced normal tissue toxicities of FLASH have now been robustly demonstrated across a number of organ systems (brain, lung, skin, gut), in a variety of animal models (mouse, rat, pig, cat, zebrafish), and by many investigators using precise dose monitoring technology [3–6]. Furthermore, the FLASH effect has been demonstrated with multiple modalities including electrons [2, 4, 6, 7], protons [8, 9], and photons [10, 11]. Feasibility has even been demonstrated in the first human clinical trial, where proton FLASH-RT resulted in tumor control and minimal normal tissue damage in patients with bone metastases [12]. FLASH-RT shows promising signs of expanding the therapeutic window and revolutionizing treatments. Although some limits and requirements to produce the FLASH-RT effect are known, the parameters that maximize normal tissue protection while providing equivalent tumor control to conventional RT are still under investigation. Key gaps in understanding persist including understanding root mechanisms, determining tissue type variability, and understanding how the FLASH effect depends on beam pulse architecture. Prior research has tended to focus on differential effects on causing DNA damage, with mixed findings [13].

Here we introduce the first experiments performed on a unique physics research linear accelerator on the campus of Duke University termed HIGS (High Intensity Gamma Source). The HIGS is of interest because of its potential for extremely high instantaneous dose rate (IDR) and deposition of doses in the range of 1 to 20 Gy in a single microsecond pulse. The HIGS is one of the accelerator facilities of Triangle Universities Nuclear Laboratory (TUNL) and can run in a customized mode capable of delivering a high-energy electron beam with FLASH-RT dose rates (we refer to this beam as the HIGS-FLASH beam). The electron beam can be extracted at two points corresponding to energies of 35 MeV or 180 MeV. A second novel aspect to the experiments reported here is the combination of HIGS-FLASH with a live-tissue *ex vivo* organotypic brain slice assay enabling more efficient, flexible, and higher-throughput investigation of FLASH mechanisms in normal brain and tumor tissue compared with *in vivo* studies. Initial beam characteristics and effects of FLASH irradiations on organotypic rat brain slices and tumor cells are presented.

## Methods

### HIGS-FLASH Linac

The HIGS linac (**Figure 1A**) is part of the larger HIGS facility on the campus of Duke University, part of TUNL [14, 15]. HIGS is a world-leading facility for nuclear physics, featuring the highest-flux Compton gamma-ray source in the world. The facility comprises an electron storage ring, a booster injector, and a linac pre-injector. The linac injects 180 MeV electrons into the booster, which raises their energy to as high as 1.2 GeV and injects the electrons into the storage ring. Electrons circulating in the storage ring emit light in undulator magnets, which is trapped inside an optical cavity to power a free-electron laser. Compton scattering of photons in the laser cavity off electrons in the storage ring produces the high-intensity gamma-ray beam at energies between 1 and 1.2 GeV. Unlike clinical linacs, the HIGS linac uses a thermionic radiofrequency (rf) gun with the rf power directly coupled into the gun cavity for immediate and rapid acceleration of the electrons after emission from the cathode (**Figure 1B**). Initial Monte-Carlo simulations of the HIGS linac suggested extremely high IDR, comparable to the highest available [16] (**Figure 1C**). We therefore utilized the electron beam from the HIGS pre-injector linac for FLASH experiments. Two positions along the HIGS linac allow extraction of the electron beam using a 9-degree dipole magnet at either 35 MeV or 180 MeV. The voltage at a Faraday cup at the beam dump enabled tracking of the dose/pulse after calibration with EBT-XD film irradiations.

**Figure 1:**
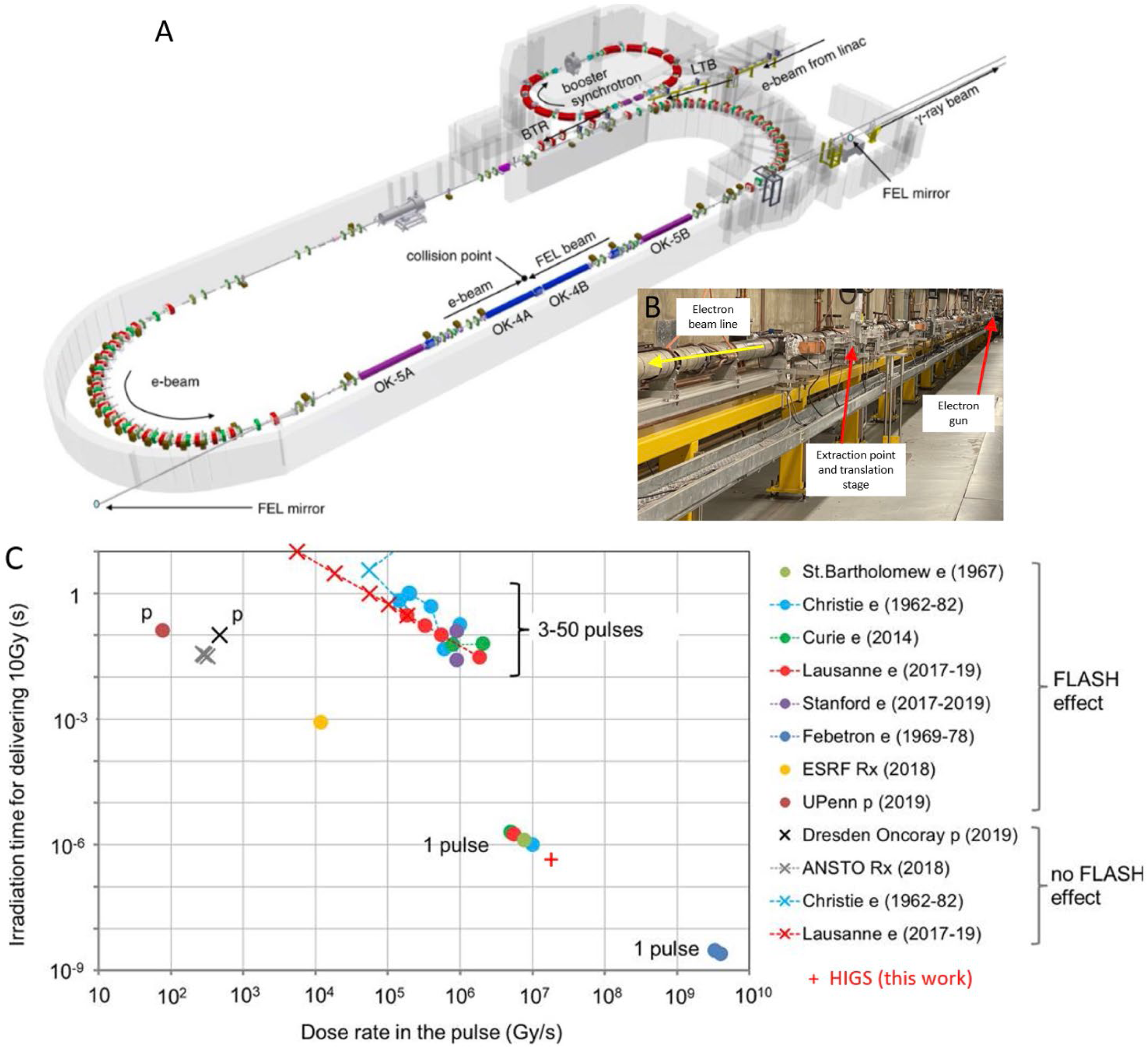
(A) Schematic of the High Intensity Gamma Source (HIGS) facility on the Duke University Campus, part of TUNL. (B) The inserted photograph shows the pre-injector electron accelerator component of the HIGS used in this work to produce a 35 MeV FLASH electron beam. Irradiation geometry at extraction point is shown in Figure 2. (C) The relation of HIGS-FLASH electron beam to other FLASH work indicating the extremely high IDR and short time to deliver total dose of 10 Gy (adapted from Montay-Gruel [36]).

### Radiochromic Film Dosimetry

Beam-spot geometry, dose rate (instantaneous and mean – IDR and MDR respectively), and well plate targeting accuracy of the HIGS-FLASH 35 MeV electron beam were determined using EBT-XD Gafchromic film dosimetry, charge measurements from the Faraday cup and pulse tracings from an integrating current transformer (ICT) at the beam extraction point. Film measurements were made by attaching pieces of film to dummy well plates held in the same orientation and position as well plates containing brain slices (**Figure 2A**). Placing film on the upstream and downstream sides (∼2.5 cm separation) enabled estimation of dose at the brain slice locations. A calibration curve, acquired by irradiating film from the same batch with 20 MeV electrons from a calibrated clinically commissioned Varian 2100ix linear accelerator (VCLA), enabled conversion of HIGS-FLASH film optical density to dose. Instantaneous dose rate estimates required timing data from the pulse shape acquired from the ICT (**Figure 2B-C**).

**Figure 2:**
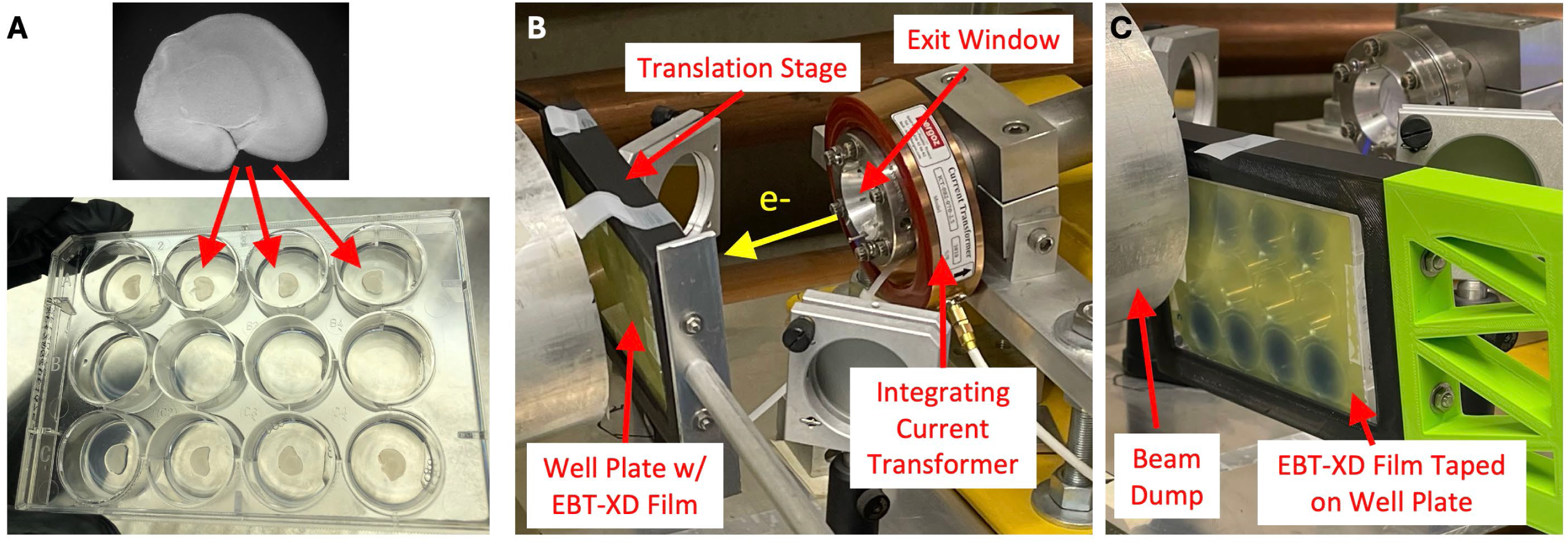
(A) a 12-well plate with 350 µm brain slices in the upper and lower rows of wells as indicated. Each slice is mounted on a ∼5 mm nutrient agarose base. (B) For irradiation, the plate is held normal to the incident electron beam at 9-degree angle to the vertical in a translation stage positioned ∼3 cm in front of the beam dump at the extraction point. (C) Close up view of the EBT-XD film attached to the rear of the well plate after a verification irradiation of 8 FLASH irradiations (4 upper row, 4 lower row). Individual irradiations of upper and lower wells are visible as darker regions on the film – the well outline is also visible on the film as lighter circles due to increased attenuation in the well walls.

### Organotypic Rat Brain Slice Model

We used an organotypic rat brain slice model to investigate the effects of FLASH irradiation on neoplastic and normal cells. This *ex vivo* model is a middle ground between *in vitro* and *in vivo* models, offering a more realistic environment for brain tumor cell growth than purely *in vitro* while generating more rapid results with lower cost than many purely *in vivo* experimental systems [17]. Preparation of the model followed methods used previously in the literature [18–20]. Briefly, brains from nine-day post-natal Sprague Dawley (Charles River) rats were divided into two hemispheres. Each hemisphere was sliced coronally with a vibratome (Leica) into 350 μm thick slices. The slices were then placed atop an agarose growth substrate in 12-well plates. As the beam shape is elliptical in cross-section, in order to minimize dose contamination from neighboring spots, the middle row of the 12-well plate was left empty, with the plate oriented such that the long axis of the elliptical beam cross-section was oriented toward the empty middle row. Each plate therefore had 8 brain slices, 4 in the upper row and 4 in the lower row, with the center row of 4 wells left empty to prevent dose overlap. The well plates were oriented such that the beam passed first through the plate lid, then through air to the brain slice, then the agarose support, then the bottom of the plate, and finally through the film before exiting to the beam dump. Film dosimetry across the depth of the well plate with simultaneous measurements from film affixed to the well plate lid and the back of the plate showed dose differences of 12% +/-6%, in accordance with the difference in distance from the HIGS electron source. The outline of the well plates is visible on the film, enabling verification of targeting accuracy (see **Figure 3A**). Care was taken to ensure that brain slices were mounted centrally in each well. To ensure targeting of each brain slice with the HIGS-FLASH beam, every HIGS irradiation was only performed after extensive alignment verification confirmed by EBT-XD film. Depending on what study was performed, brain slices were seeded with 4T1 mouse breast carcinoma cells expressing an mCherry/firefly luciferase dual reporter construct [20] or normal neurons in the brain slices were biolistically transfected with plasmid encoding yellow fluorescent protein (YFP) [21]. Normal tissue studies of cytokine effects and confocal microscopy used the brain slice alone without introduction of YFP transfection or cancer cells.

**Figure 3:**
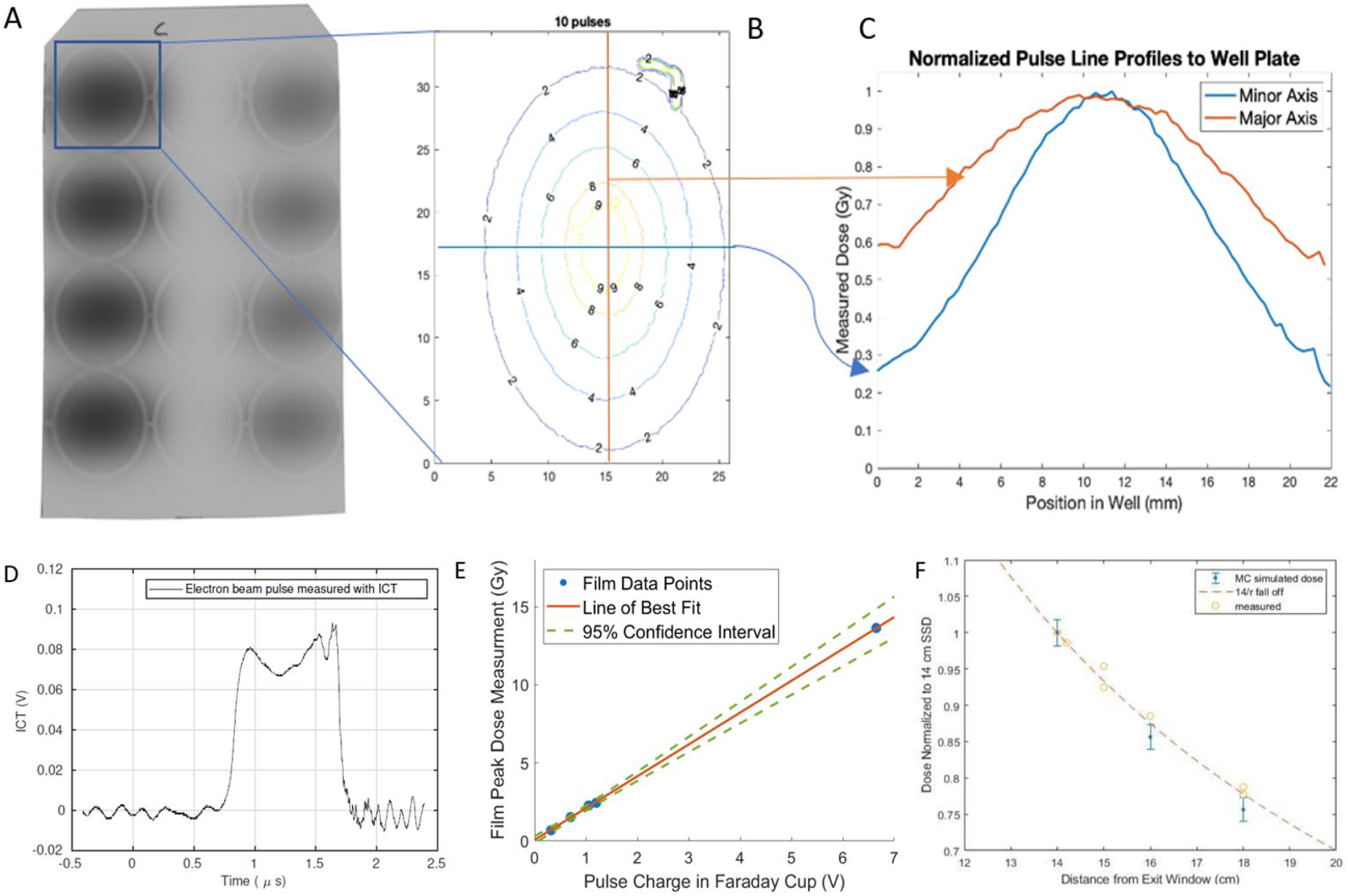
(A) Scanned image of EBT-XD film taped to the back of a well plate after HIGS-FLASH irradiation of all wells in the upper and lower rows (note the film is shown here rotated 90 degrees – its actual orientation during irradiation is shown in Figure 2). The middle row was unused to avoid dose contamination from neighboring beams. (B) An isodose plot of the HIGS linac pulse distribution normalized to the peak dose. (C) Line profile through the ellipse major and minor axes showing near Gaussian dose distribution along both the major and minor axis. (D) Timing data acquired from the integrating current transformer (ICT shown in Figure 2) for a single 15 Gy pulse from HIGS-FLASH beam. (E) Linear dependence of the voltage on the Faraday cup with peak dose delivered to the brain slices. (F) Variation in dose as the plates are moved away from the exit window illustrating ∼ 1/r falloff.

### Cancer Cell Control Studies

Rat brain slices were prepared as above for culture on agarose substrate in 12-well plates and were seeded with mCherry-firefly luciferase-expressing 4T1 murine breast cancer cells at a density of 4000 cells per brain slice. Plates were irradiated either with the HIGS 35 MeV electron beam at varying dose rates, or with a 20 MeV conventional electron beam from the VCLA. For HIGS irradiations, 8 slices per plate were seeded with the middle row left empty to avoid dose contamination from irradiations of neighboring wells. HIGS irradiations were performed with different pulse sequencing and amplitudes to explore the effect of different MDR (mean dose rate) within the total dose range, yielding a measurable dose response of the 4T1 cell line (0 – 7.5 Gy). The intra-pulse IDR for all HIGS irradiations was very high, ranging 2 – 6 MGy/s. HIGS-FLASH irradiation consisted of a single pulse delivering nominally 6 Gy in 1 µs gave an estimated MDR of 6.17 x 10^6^ Gy/s. HIGS irradiations at other MDR and were delivered in multiple pulses with a 0.4 s repetition time to total doses of 2.5 Gy (2 pulses), 5 Gy (4 pulses) and 7.5 Gy (6 pulses) at MDR rates between 3 – 6.25 Gy/s. During each HIGS irradiation, the well plate was suspended as shown in **Figure 2B,C**. Prior to irradiating a plate containing brain slices, accurate set up and targeting was verified on a duplicate dummy test-plate containing EBT-XD film. Test irradiations were visually inspected to ensure accurate targeting of each individual well in the upper and lower rows of the plate. The mCherry and firefly luciferase imaging were performed 4 - 5 days after irradiation as described below.

For conventional VCLA irradiations, we replicated the doses delivered in HIGS irradiations using a Varian 2100EX clinical linac (VCLA) operating at 1000 MU/min with 20 MeV electrons. 12 brain slices per plate were used for conventional VCLA irradiation where all wells were irradiated at once. Plates were positioned on top of 9 cm solid water on the treatment couch and irradiated with a 20 x 20 cm AP field. The dose rate at the brain slices for all VCLA irradiations was 0.17 Gy/s, well below the FLASH range. Unirradiated control plates of brain slices were included with each experiment and undertook the same handling, transport, temporal, and storage histories.

### Cancer Cell Imaging and Analysis

mCherry fluorescence was monitored with live-cell imaging following the methods outlined in [22]. Briefly, mCherry fluorescence and brightfield images of the live brain slices were obtained immediately before irradiation, and daily afterwards with a Zeiss Lumar V12 stereoscope. Stereoscope images were analyzed using CellProfiler software (https://cellprofiler.org) to identify mCherry-positive cells and determine integrated fluorescence intensity as a proxy for cell growth. Customized image analysis pipelines are available on request, while further details are described below and in the Supplemental Material and in [16, 22]). On day 5 post-irradiation, luminol reagent (XenoLight D-Luciferin, Perkin-Elmer Cat.# 122799) was added to the wells on top of the brain slices before incubation for 10 minutes at 25 degrees C, and firefly luciferase signal was measured with a Lumina XR IVIS bioluminescence imager (Perkin-Elmer) following the methods used in [20].

### Normal Tissue Studies

350 µm-thick brain slices were prepared as described above. For neuron morphology assessments, the methods were adapted from established studies of stroke and neurodegenerative disease [19,23,24]. Brain slices were transfected immediately after slicing with YFP plasmid (Gwiz expression vector, Genlantis) using a BioRad Helios gene gun and 1.6 mM gold particles. All slices on a plate were irradiated with either the HIGS-FLASH beam or the conventional VCLA 20 MeV electron beam operating at 1000 MU/min (irradiation set up and technique was the same as the cancer cell study described above). Unirradiated plates that followed the same handling, transport, temporal, and storage histories served as controls. The HIGS irradiations were carried out under FLASH (0.15 seconds inter-pulse) or sub-FLASH (10 seconds inter-pulse) conditions. HIGS-FLASH irradiations were typically 5 - 6 Gy per pulse. EBT-XD film was used to determine the total dose and dose rate delivered to each slice. Three independent normal tissue health assays were carried out: 1) YFP fluorescence and neuron morphology were scored daily by live-cell stereoscope microscopy out to 5 days; further details are given in supplementary **SFigure 1**. 2) Slice washes on day 3 post-irradiation (200 µL phosphate-buffered Ca/Mg-free saline incubated with slices for 30 min at 37C and 5% CO2) were analyzed for cytokine expression with MILLIPLEX® Rat Cytokine/Chemokine Magnetic Bead Panel (Millipore-Sigma) processed at Eve Technologies, Calgary, AB, CA (RD27 assay). 3) Slices were fixed in 4% paraformaldehyde on day 3 post-irradiation and processed for immunofluorescence microscopy with a Leica SP5 scanning confocal microscope Thermo Cat.# D1306), IBA-1 for microglia (FUJI Film Wako Chemicals Cat.# 019-19741) and GFAP for astrocytes (Sigma-Cat.# MAD360). Confocal images were analyzed and processed with customized in-house pipelines developed in CellProfiler (https://cellprofiler.org). Further details can be found in the Supplemental Material **SFigure 2**. Significance comparisons were assessed by One-way Brown-Forsythe and Welch ANOVA tests performed using GraphPad Prism 10.

## Results

### Characterizing the HIGS-FLASH Beam

We first examined beam-spot geometry and well plate targeting accuracy using EBT-XD radiochromic film dosimetry. The EBT-XD spot-size of the HIGS-FLASH beam is shown in **Figure 3A**. The penumbra of the beam spot (**Figure 3B**) was slightly wider in the vertical orientation (major axis in **Figure 3C**). 80% of the peak dose is received by an elliptical region of dimension 11 x 7 mm. This area covers the entire rat brain slice provided the HIGS-FLASH beam is accurately placed centrally on the surface, enabling the brain slice studies below.

The dose rates of the HIGS-FLASH were determined from EBT-XD calibrated film measurements. The timing data for a single pulse is shown in **Figure 3D** yielding a measured instantaneous dose rate (IDR) of the HIGS 35 MeV electron beam as up to 20.7 ± 0.6 MGy/s at the plane of the brain slices. The linear dependency of the size (Gy) of HIGS pulses versus voltage applied to the HIGS cathode is shown in **Figure 3E**. Lower IDR can easily be selected by lowering the voltage on the cathode as per this relation which was observed to extend up to maximum voltages of 6.6 V and maximum pulse size (so far) of 20.7 Gy. We characterized dose falloff with distance from the exit window at extraction point as shown in **Figure 3F**. Thus, the HIGS-FLASH beam provides a tunable high energy electron FLASH beam. Further, the HIGS-FLASH can deliver one of the highest intra-pulse mean dose rates of modern FLASH beams.

### HIGS-FLASH Effects in Neoplastic Cells

A key element of clinically useful FLASH-RT is that while normal tissues are spared, tumor cells are not and can still be controlled by FLASH radiation, resulting in a widened therapeutic window. To test this, we employed an organotypic brain slice model of breast cancer brain metastasis [22,25]. The results of irradiating 4T1 murine breast cancer cells (labeled with both mCherry and luciferase) in the brain slice model are shown in **Figure 4**. These data were acquired 4 - 5 days after irradiation.

**Figure 4:**
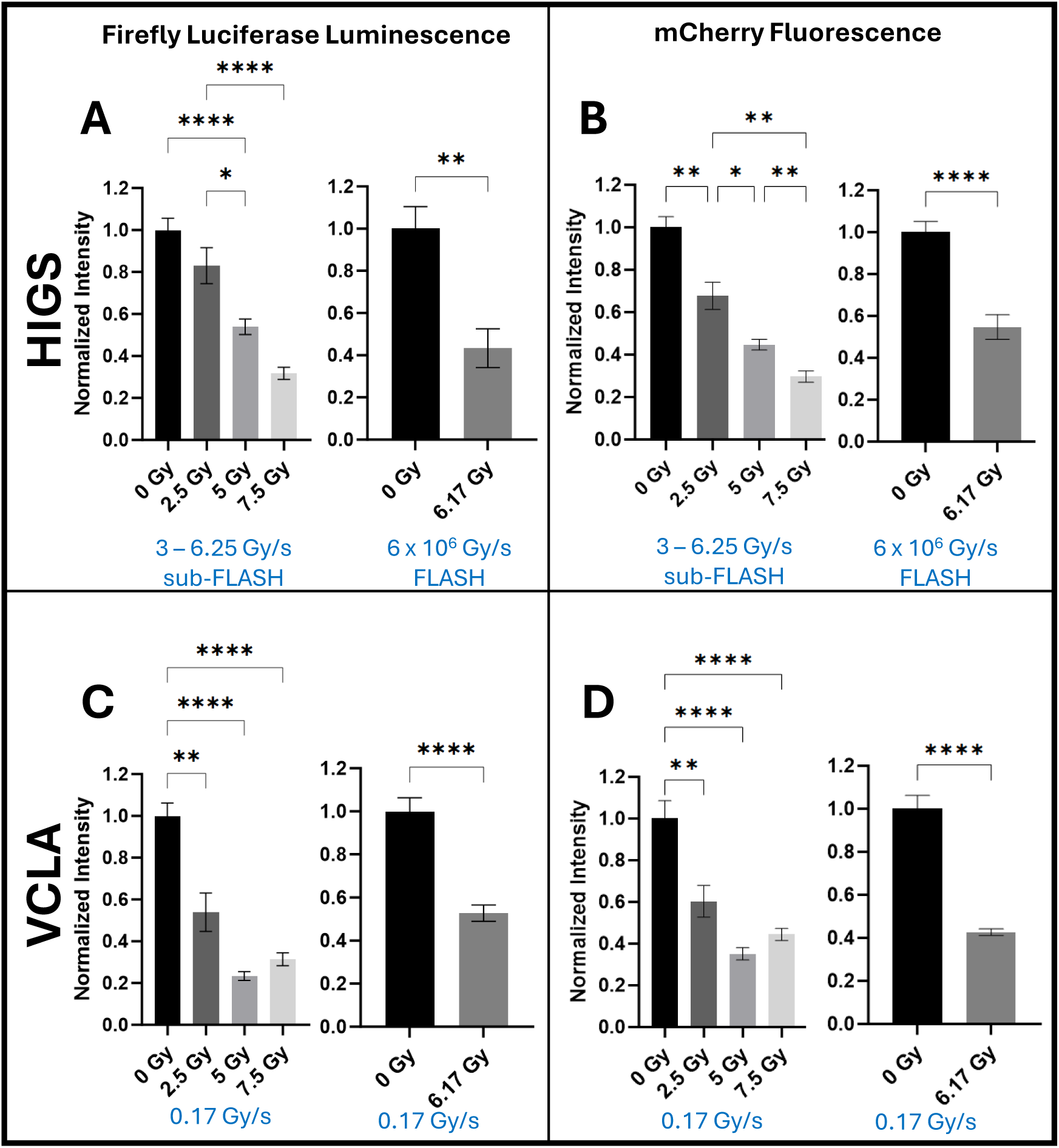
Rat brain slices seeded with 4T1 mCherry FLuc cells were irradiated at either the HIGS (A,B upper row) or the Varian clinical linear accelerator (VCLA). (C,D lower row). Brain slices were then analyzed for tumor burden by two independent assays: firefly luciferase (left column) and mCherry fluorescence (right column). The dose rate for all VCLA irradiations was 0.17 Gy/s. On the HIGS, selecting different pulse amplitude and sequencing enabled irradiations with different MDR. The IDR for all HIGS irradiations was very high (range 3 - 6.17 x 10^6^ Gy/s). The HIGS sub-FLASH MDR irradiations were delivered in multiple pulses, each separated by 0.4 seconds to total doses of 2.5 Gy (2 pulses), 5 Gy (4 pulses) and 7.5 Gy (6 pulses). The MDR for this group was in the range of 3 – 6.25 Gy/s, sub-FLASH per the 40 Gy/s rule. The HIGS FLASH irradiation consisted of a single pulse delivering ∼6 Gy instantaneously giving a MDR of ∼6 x 10^6^ Gy/s. The mCherry signal was assessed once daily via stereoscope images for 4 - 5 days after irradiation, and CellProfiler segmented the fluorescent signal to give a measure of total intensity. The terminal firefly luciferase assay was performed at final endpoint. Mean and S.E.M are shown for n ≥ 8 biological replicates. Each assay was tested for significance using a Brown-Forsythe and Welch ANOVA test (for experiments involving three or more cohorts) or a t-test (for experiments involving two cohorts). All assays have at least two groups with significant difference in means where p < 0.0001. The time plot of the mCherry data is given in supplementary material **SFigure 3**. An additional plot of the above data arranged for 1:1 comparison can be found in **SFigure 4**.

A strong dose response was observed using independent mCherry fluorescence-based and luciferase-based assays with both FLASH (**Figures 4A,B**) and conventional VCLA beams (**Figures 4C,D**). Unirradiated control slices demonstrated exponential growth of 4T1 cancer cells (see supplementary material **SFigure 3**). These observations are consistent between both the quantified stereoscope live-cell images (**4A, 4C**) and the independent luciferase analysis (**4B, 4D**), indicating comparable isoeffective growth inhibition for FLASH, sub-FLASH, and VCLA dose rates.

### HIGS-FLASH Normal Tissue Effects

#### Cytokine Analysis

FLASH-RT, compared to conventional RT, has been reported to reduce astrogliosis and brain inflammatory changes as early as two weeks post-irradiation [26]. Early differences in radiation-triggered release of secreted inflammatory signaling molecules could underlie these observations. To investigate this potential, we turned to profiling cytokines released from the brain slices following irradiation at different dose rates. Intriguingly, as shown in **Figure 5**, elevated levels of TNFα and especially fractalkine were released following irradiation with FLASH dose rates.

**Figure 5:**
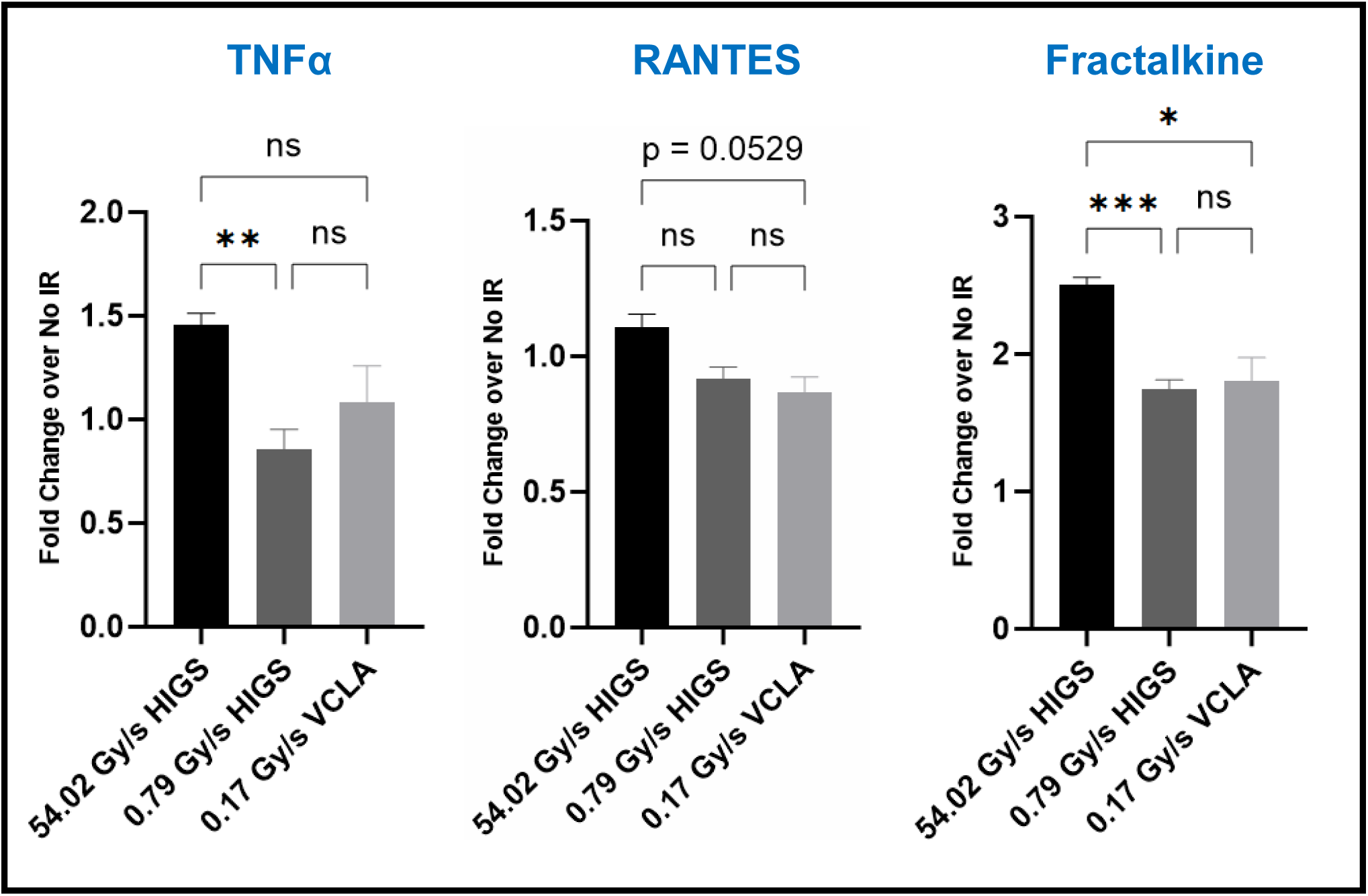
Profile analysis of cytokines released from brain slices irradiated at the indicated dose rates with either a conventional clinical linear accelerator (0.17 Gy/s VCLA) or using the HIGS-FLASH beam line (0.79 Gy/s and 54.02 Gy/s). Measured total irradiation doses were ∼24.31 Gy for all samples. Cytokine levels were normalized to unirradiated controls and expressed as fold change. Mean and S.E.M. are shown for n ≥ 4 biological replicates. Significance was determined by a One-way ANOVA test with Brown-Forsythe correction.

RANTES, though not significantly upregulated by FLASH, demonstrated a suggestive trend of increased expression at higher dose rates with p = 0.0529. The elevated release of these cytokine mediators indicates that immune mechanisms may underlie early responses to ultra-high dose rate irradiation of normal brain tissue. A heat map of the complete cytokine profiling is shown in supplementary material **SFigure 5**.

#### Microglial Analysis

We next investigated microglial characteristics following irradiation at differing dose rates, as fractalkine has been implicated in activation of microglia [27]. Using confocal microscopy of slices fixed 3 days following irradiation, we observed striking differences in the appearance and morphology of microglia (revealed by IBA-1 staining) following irradiation at FLASH dose rates (illustrative results in **Figure 6A-D**). These differences include enlarged size of microglial cells and an increased stellate morphology at a FLASH dose rate of ∼54 Gy/s. Area (**Figure 6E**), perimeter (**Figure 6F**), and mean radius (**Figure 6G**) of microglia in HIGS-FLASH irradiated slices were larger than microglia irradiated at a sub-FLASH dose rate, reaching high statistical significance (p < 0.0001).

**Figure 6:**
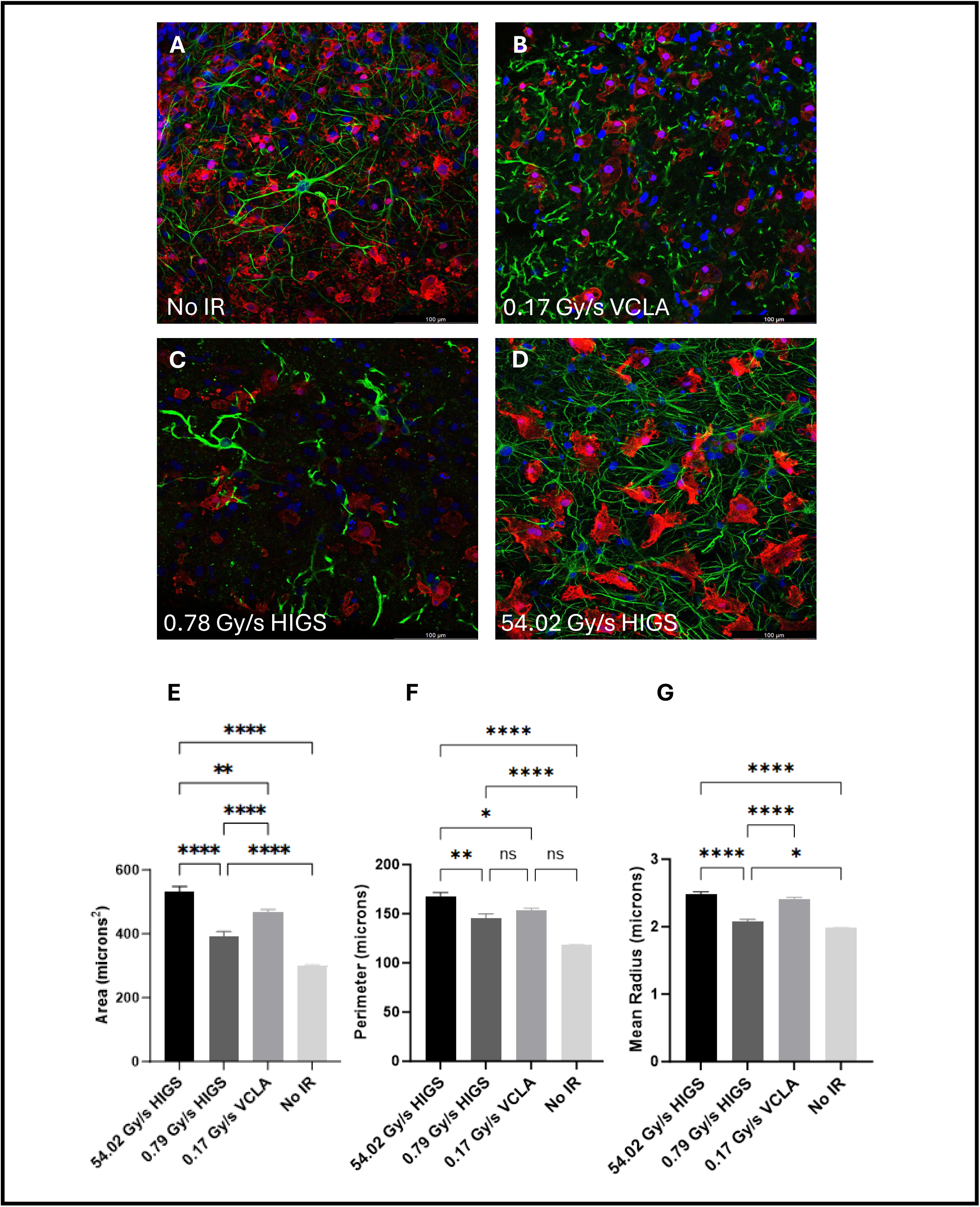
(A-E): Confocal images of rat brain slices fixed and stained 3 days post-irradiation with ∼24.31 Gy. Red: IBA-1 (microglia); Green: GFAP (astrocytes); and Blue: DAPI (nuclei). (A) unirradiated control, (B) VCLA MDR 0.17 Gy/s, (C) HIGS sub-FLASH MDR 0.79 Gy/s, and (D) HIGS FLASH MDR 54.02 Gy/s. (E-G) CellProfiler analysis of fluorescent confocal images for indicated measures of microglial morphology. Mean and S.E.M. are shown for n=15-23 images per condition. Significance of morphology signal segmentation was tested for using a Brown-Forsythe and Welch ANOVA test, and **** denotes p < 0.0001.

#### Neuron Morphology Analysis

Current evidence indicates that FLASH irradiation alters early microglial responses to RT without immediately impacting neuronal health [26]. We therefore performed neuron morphology assays to test if HIGS-FLASH irradiation had acute effects on neuronal health in the brain slice assay. These experiments showed that neuron morphology and viability was generally unaffected by irradiations in this dose range, regardless of dose rate. Illustrative stereoscope live-cell images of a brain slice imaged on 3 consecutive days are shown in supplemental **SFigure 2**. A trend was observed in that the number of healthy-appearing neurons decreases with each passing day, as expected in this transient transfection system and short-term brain slice culture conditions.

## Discussion

Here we characterize HIGS-FLASH, a unique high energy electron beam that provides a tunable FLASH experimental setup with extremely high IDR and MDR capability. The HIGS-FLASH 80% isodose line creates an elliptical 11 x 7 mm beam spot, large enough to effectively treat a hemi-coronal p9 rat brain slice as described, and would also encompass a whole mouse brain *in vivo*. We linked the HIGS-FLASH accelerator to a flexible rat brain slice model system and observed that FLASH dose rates (as compared to sub-FLASH or VCLA dose rates) had comparable effects on 4T1 cancer cell growth inhibition, and differential effects on acute cytokine release and microglia morphology, while normal neuronal morphology was largely preserved.

In comparing effects of our HIGS linac to other irradiation sources, there are some unavoidable differences between clinical linear accelerators and the HIGS linac. One of the differences between accelerators is in electron energy. While we can extract the HIGS electron beam at either 35 or 180 MeV, we chose to study 35 MeV, as this is closest to the highest electron energy available to us from clinical treatment machines, 20 MeV. EBT-XD dose measurements at 35 MeV and 20 MeV were directly compared (i.e. no energy correction was necessary) because over this energy range the film sensitivity does not vary appreciably and RBE ≈ 1.0 [28]. Given the 350 µm brain slice thickness in our cancer cell assay and the similarity in relative biological effectiveness and surface characteristics for electrons at 20 MeV and 35 MeV [28], this difference in electron energies is unlikely to be important. We also chose to vary the mean dose rate on the HIGS linac itself to compare effects of dose rate more directly. Observations on cancer cell growth inhibition and normal neuron morphology were comparable across dose rates and different linacs. Interestingly, we observed differential effects of FLASH dose rate, consistent across linacs, on immune cells and signaling molecules in the brain slices.

The cytokine and microglia findings illustrate the acute effects of this HIGS-FLASH beam on the rat brain slice model. In our survey of cytokine responses, effects on fractalkine were most pronounced, and HIGS-FLASH irradiation induced higher fractalkine fold change compared to all other dose rates. Other studies report that in irradiated mouse microglia, fractalkine can change microglia from the pro-inflammatory M1 state into the anti-inflammatory M2 state [27,29]. Furthermore, that same study found that higher levels of fractalkine decreased memory loss for mice irradiated cranially with 10 Gy. However, the relationship of fractalkine signaling to long-term changes in microglia and ultimately brain function is complex, with other reports indicating that fractalkine can induce pro-inflammatory changes, and that the timecourse of acute and late effects of fractalkine may differ [30,31]. In addition to fractalkine, TNFα also increased after FLASH irradiation compared to other dose rates. TNFα is reported to be a pro-inflammatory cytokine that is expressed by microglia in the M1 state [32]. For RANTES, a cytokine that directs immune cells towards inflammation [33], our data suggest a strong trend that HIGS-FLASH irradiation also increases RANTES release. Taken together, these data suggest that the ultra-high dose rates achieved by the HIGS-FLASH accelerator induce different cytokine profiles compared to low dose rates. These differences in cytokine release could have important consequences for both radiation-induced neuroinflammation and cognitive function, and may also lead to novel brain tumor therapies [34].

Based on these findings of differences in cytokine release in brain slices following HIGS-FLASH irradiation and their links to microglia, we investigated microglial morphology in the brain slices. HIGS-FLASH irradiation caused significant alterations in several features of microglial morphology. Although the full picture of more long-term changes in microglia and brain inflammatory processes is not yet clear, certainly these acute changes induced by ultra-high dose rate electron irradiation merit further investigation.

It is notable that other studies have identified inflammatory responses in the brain that involve microglia as part of the normal tissue sparing effect observed in the brain at FLASH dose rates [35,36], although there are key differences with the current study. We point out that our brain slice system uses *ex vivo* culture of brain slices from post-natal day 9 rats. Our findings in juvenile rat tissue that has undergone a traumatic insult in the slicing process could differ from studies of intact adult mouse brains. Nonetheless, the identification of acute cytokine and microglial changes induced by ultra-high dose electrons in our model system does point to the involvement of neuroinflammation in general as an important pathway in the mechanism of normal tissue effect of FLASH irradiation. It is also tantalizing to speculate about possible therapeutic implications of inflammatory changes induced in microglia and macrophages by HIGS-FLASH dose rates given intriguing recent data obtained with proton FLASH irradiation in mouse brain tumor models. The brain slice assay presented here provides a rapid and low-cost tool to explore these early responses.

Paired with the HIGS-FLASH beam that is capable of mean dose rates ranging from ∼0.2 Gy to 20.7 MGy/s, we are hopeful that the effects and mechanisms of dose rate on radiation-induced brain inflammation can be further elucidated. The HIGS-FLASH beam will enable investigations into effects not only in the current accepted “FLASH” dose rate range (greater than 40 Gy/s), but beyond into extremely high dose rates of MGy/s. We anticipate that additional insights into mechanisms of FLASH effects on normal tissues can be elucidated by extending dose rates to this range. Looking forward, it is clear that the HIGS-FLASH beam line has certain challenges for translation to clinical use. As currently configured, treatment of human patients would be limited to very small tumors given the small field size. The very high energy (180 MeV) capability of the beam line is intriguing, as this could translate to external beam treatment of deep tumors. We note that there are commercial enterprises who are developing similar technologies with very high energy electrons, but with larger field size capabilities. To our knowledge, these technologies are still in the planning stages and we are hopeful that the HIGS-FLASH beam could help elucidate FLASH mechanisms and perform pre-clinical testing that could help guide the development of FLASH electron therapy for human patients.

There are some limitations with this study. One limitation involves the brain slice model used to investigate the normal tissue and cancer cell effects of the HIGS-FLASH beam. While this model captures more biological complexities than *in vitro* studies like the tumor microenvironment through 3-dimensional cell-cell contacts, it does not completely capture the biological complexities of *in vivo* models [25]. Furthermore, the short lifespan of the model limits the study of the model to acute responses rather than long-term effects. Another limitation is the confocal imaging of the microglia. These images provide information on a 2D slice of the microglia, but do not fully capture 3D morphology.

## Conclusion

The HIGS-FLASH and brain slice experimental platform is an efficient and powerful new tool for mechanistic research, with extremely high IDR and MDR capability (∼20 MGy/s for both IDR and MDR with a single pulse of up to ∼20 Gy delivered in 1 µs). Initial results indicate that the HIGS-FLASH beam is effective at inducing growth arrest in cancer cells and induces acute differential cytokine and microglial effects in the brain. Our results point to a role for neuroinflammation in the acute phase of normal brain responses to ultra-high dose rate electron radiation and are consistent with other reports suggesting a fundamental role for brain inflammatory processes in the underlying mechanism of normal brain and neurocognitive function preservation following irradiation at ultra-high dose rates.

## Supplementary Material

### Confocal image acquisition and analysis

Brain slices were stained for microglia and GFAP as described in Methods and imaged with a Leica SP5 confocal microscope. The laser power settings were set to 5% for the DAPI 405 nm laser, 13% for the HeNe 633 nm laser, and 18% for the Argon 488 nm laser. A 40x oil immersion objective and zoom of 1x were chosen for imaging. The gain for each laser was set to 750V, 700V, and 800V for the DAPI 405 nm, HeNe 633 nm, and Argon 488 nm lasers respectively. All these settings were kept consistent over the multiple imaging sessions. 2D images were taken of multiple slices, with raw individual channel images exported for analysis.

All confocal images acquired in this study were analyzed and processed with customized in-house pipelines developed in CellProfiler (https://cellprofiler.org). These pipelines have been developed over several years and rigorously tested for accuracy and uncertainty, of which further details can be found in [16, 22], and pipelines can be provided on request. Care was taken to manually check the output at regular intervals in the sequence for each individual application. The basic steps are illustrated in supplementary data **SFigure 1** for the pipeline used to identify and parametrize microglial cells and astrocytes.

**SFigure 1:**
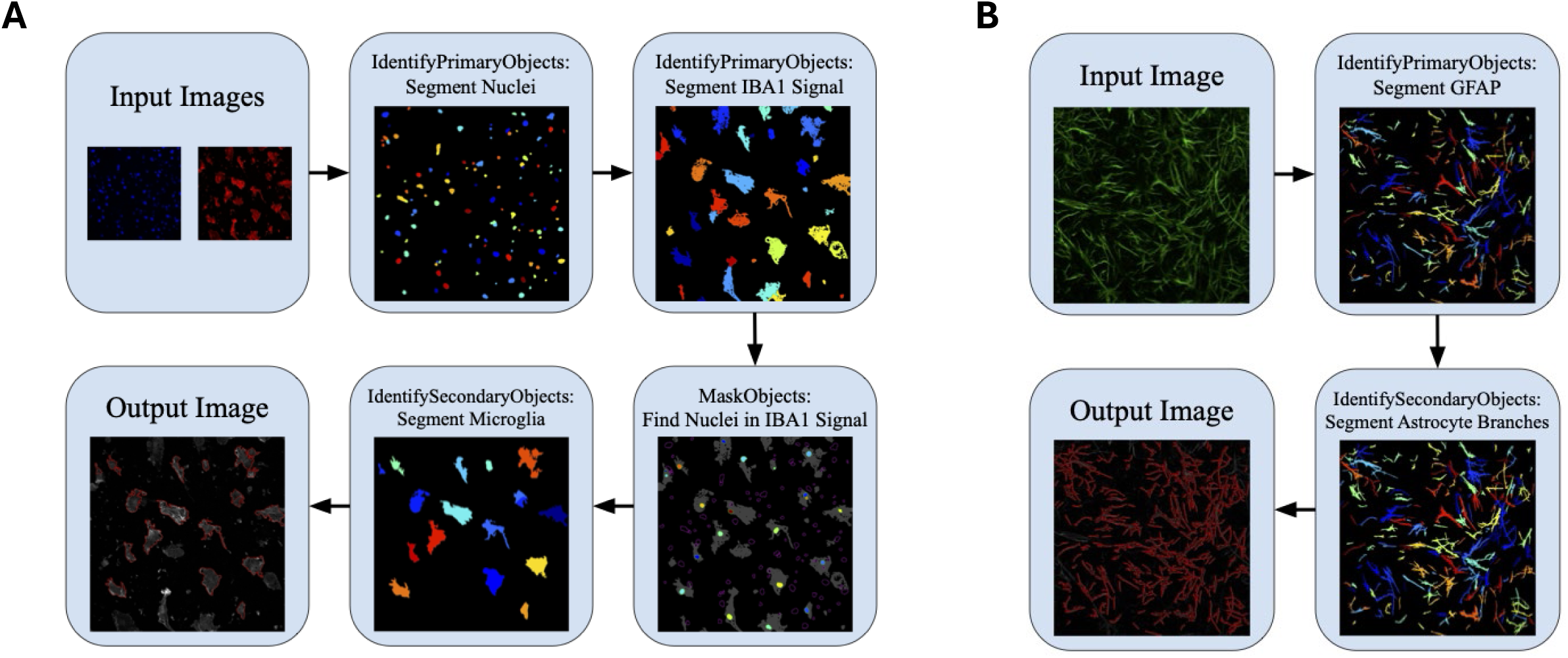
Representative real-data output from CellProfiler pipelines to analyze confocal images for (A) microglial cells, and (B) astrocytes. Color coding indicates distinct objects identified during different steps of the pipeline processing.

Morphological assessment was performed by manual inspection of YFP labelled brain slice neurons. This assay was adapted from neurodegenerative disease and stroke experiments. In these assays, variations in neuron health are measured by evaluating the morphology of the soma, axon and dendritic tree using neuron-specific fluorescent protein expression and live cell microscopy with a fluorescent stereoscope and direct observation [19,23,24]. Slices were imaged daily for five days post-irradiation using a Zeiss Lumar V.12 stereoscope. Healthy neurons were manually tracked to determine surviving fractions and YFP intensity. As shown in supplementary **SFigure 2**, the YFP labelled neuron assessment technique used in this work was as follows: the day one through day five images for a slice were loaded into FIJI as an image sequence, and the brightness and contrast were kept constant throughout the process. The only time the display settings were changed was if a given neuron was difficult to discern from adjacent neurons. After the neurons were discerned from one another, the display settings were returned to their original setting. Each visible healthy neuron on day one post-irradiation was marked using the marker feature in FIJI. After all the visible healthy neurons in the slice were marked, the markers were saved as an ROI, and the total number of markers was recorded. These markers were then kept as the image was switched between the day one and day two images. By switching back and forth between the day one and day two images, the day one healthy neurons could be found in the day two image. The shape of the neuron and its nearby surroundings were used to ensure that the correct neuron from day one was found in the day two image. The health of this neuron on day two was then assessed; if the neuron showed signs of axon breakage or lost its axon altogether, it was considered unhealthy, was not scored, and its marker was removed. If the neuron continued to be healthy, then it was given a day two marker, and its day one marker was removed. This continued until all day one markers were removed and all day two markers were placed. The day two markers were all saved as an ROI and the total number of markers recorded. This process was then repeated for each day until the day five markers were placed and recorded. The scores for each day were then normalized to day one. Finally, this process was repeated for each slice for each treatment arm. Once all the surviving fractions were determined, the mean surviving fraction was calculated for each day for each treatment arm and plotted with 95% confidence intervals and intra-reader variability added in quadrature.

**SFigure 2:**
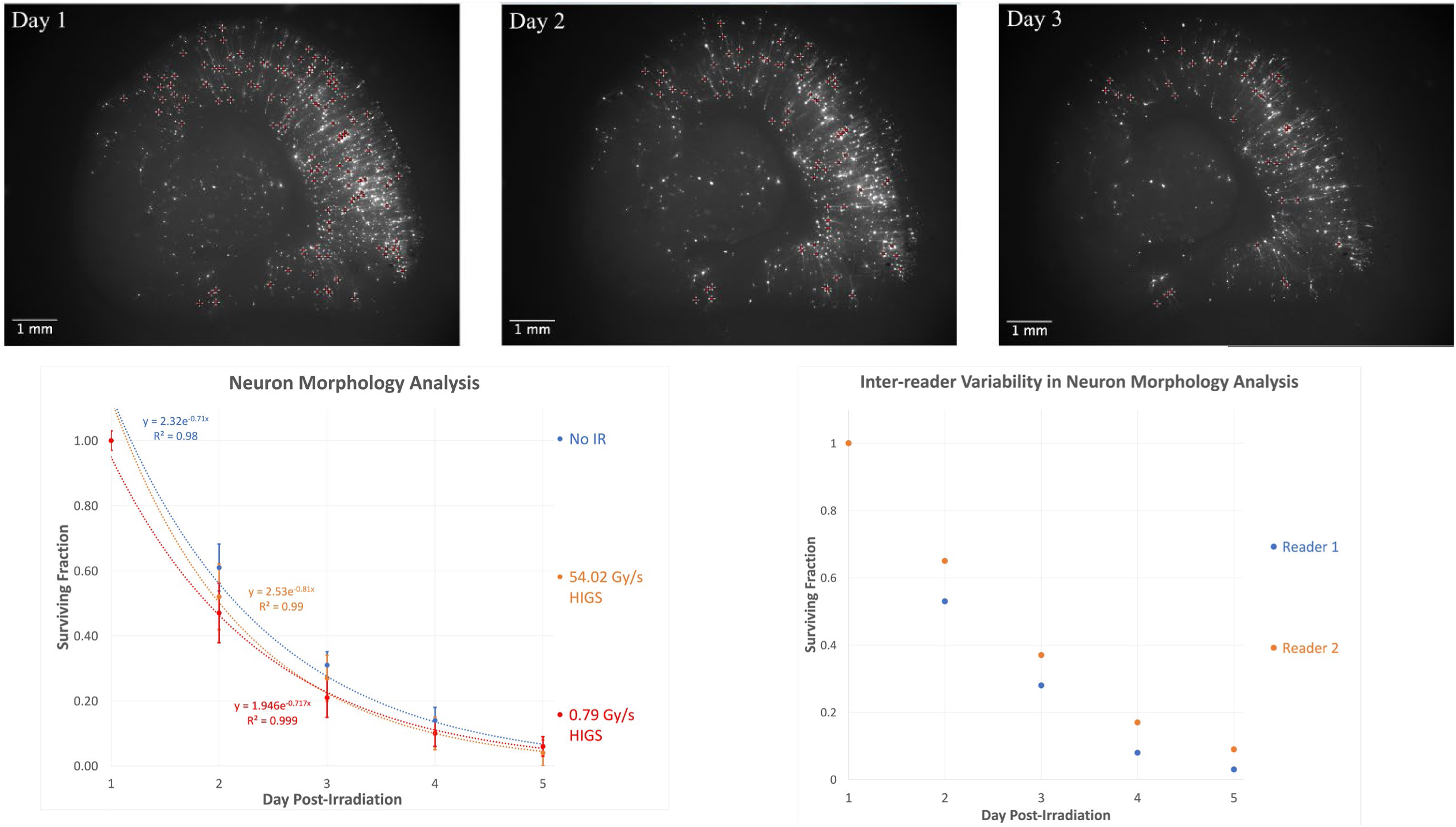
Upper row illustrative stereoscope images of a brain slice indicating manually localized healthy neurons. Lower row images (left): average healthy neuron scores over 7 slices irradiated at 54.02 Gy/s, 6 slices irradiated at 0.79 Gy/s, and 8 control (unirradiated) brain slices. (Right) An inter-observer study showed consistent trends but indicated an offset uncertainty in the absolute determination of what constitutes a healthy neuron.

Longitudinal live-cell imaging analysis of fluorescently labeled tumor cells was completed to assess the cytotoxic efficacy of the HIGS-FLASH beam compared to that of a Varian 2100ix clinical linear accelerator (VCLA) (first row, panels A,B). This was performed by assessing the total integrated intensity of the mCherry signal from episodic images taken of the 4T1 mCherry FLuc cells seeded on the P9-P11 350 µm hemi-coronal Sprague Dawley rat brain slices. Brain slices and associated cells were imaged daily for five days (both pre- and post-irradiation) with standard conditions using a Zeiss Lumar V.12 stereoscope and 1500 ms exposure with a Rhodamine filter. Image zoom and framing were kept constant for each day. These images were then exported as raw-signal .tiff files and fed through our validated CellProfiler program to segment areas of mCherry signal. CellProfiler data export of the integrated intensity of the mCherry signal was then averaged on a per-condition, per-day basis to create a longitudinal representation of cell growth over the course of the experiment. For each condition, the average total integrated intensity was expressed using the standard error of the mean as a single data point associated with the time the image was taken. At endpoint Day 5, significant differences in mCherry signal were measured using an Ordinary One-way ANOVA.

**SFigure 3:**
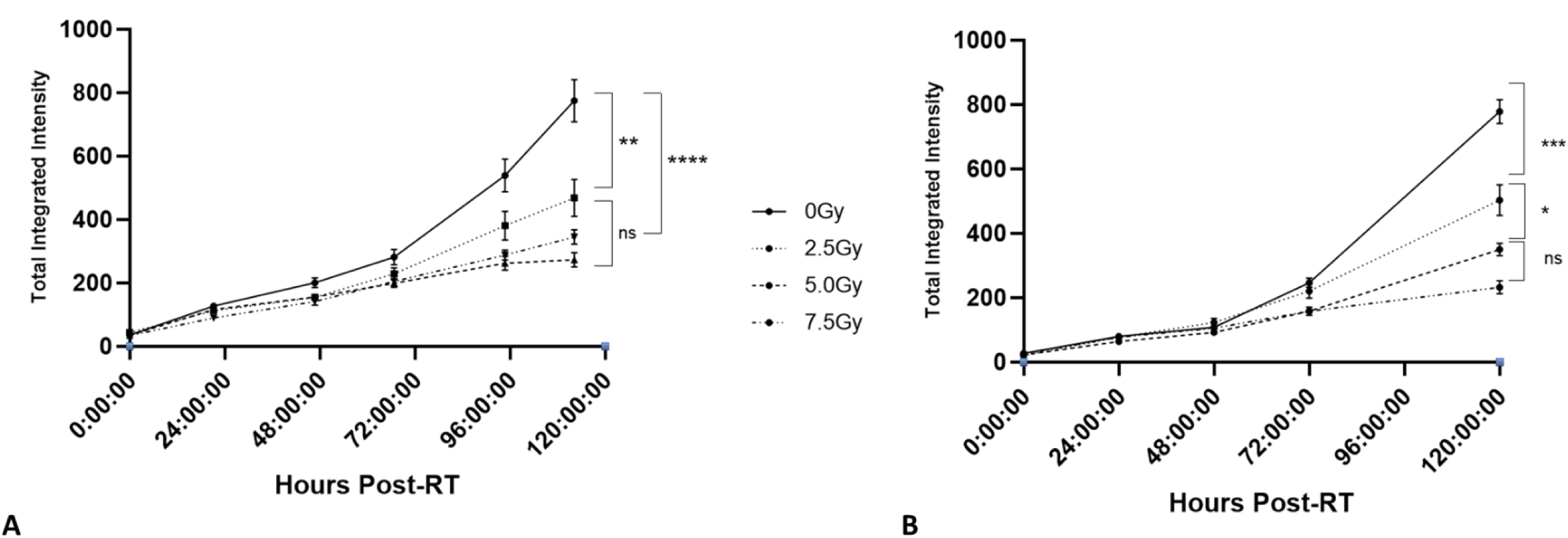
Graphs representing daily 4T1 mCherry FLuc cell growth as expressed by CellProfiler-segmented mCherry signal using different dose rates of 2.5, 5.0, and 7.5 Gy. (A) Data representing longitudinal cell growth following irradiation using the VCLA with MDR 0.17 Gy/s. Increase in dose correlates with decrease in positive slope of growth, indicating a dose-dependent viability response of treated cells. (B) Data representing longitudinal cell growth following irradiation using the novel HIGS beam with MDR 3 – 6.25 Gy/s. The same dose-dependent viability trend found in the conventional dose rate experiment can be seen, which indicates that the HIGS-FLASH beam is capable of comparable tumor control to a conventional clinical system.

**SFigure 4:**
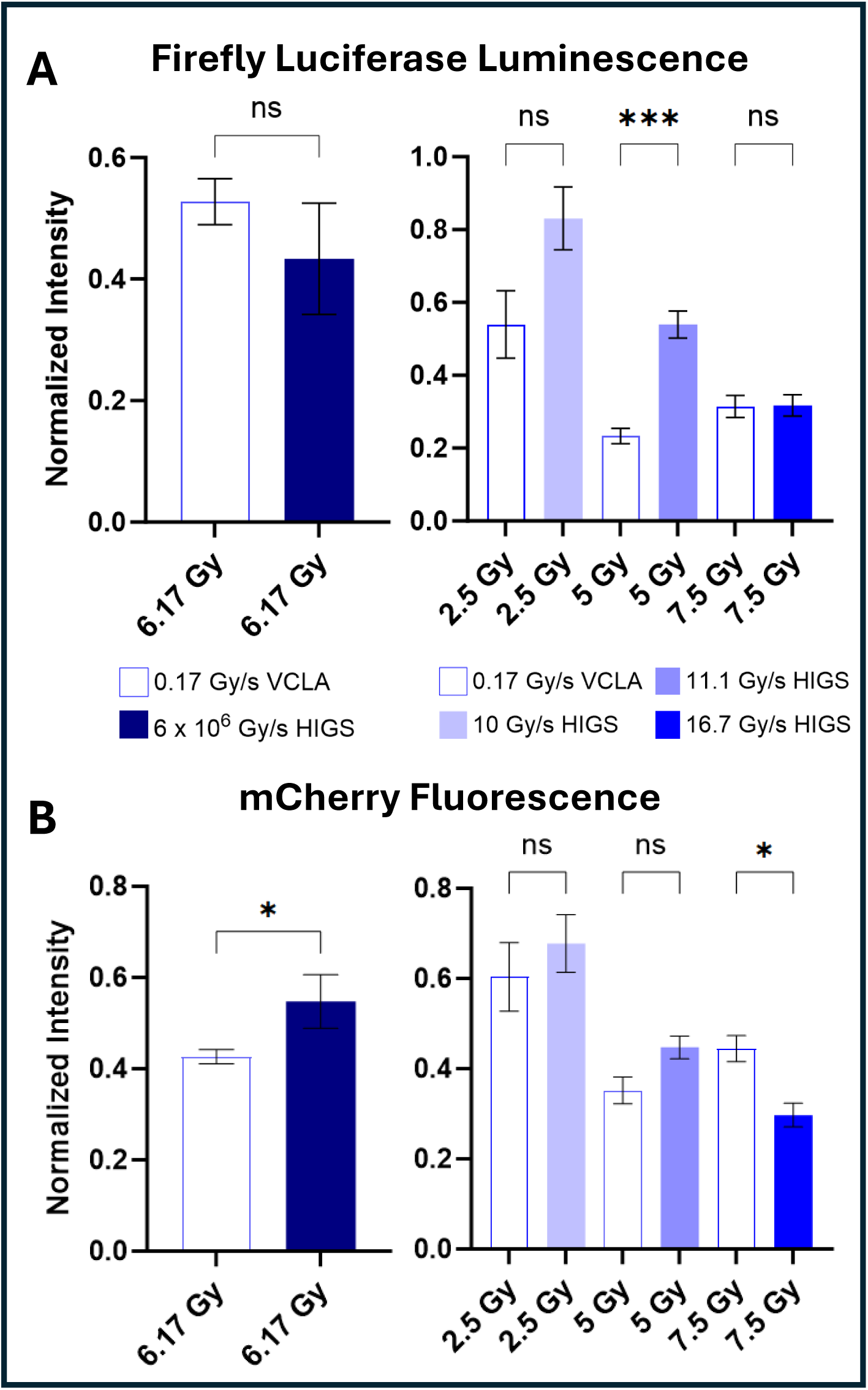
Rat brain slices seeded with 4T1 mCherry FLuc cells were irradiated at either the HIGS or the Varian clinical linear accelerator (VCLA) for doses ranging from 2.5 - 7.5 Gy at varying dose rates. Brain slices were then analyzed for tumor burden by two independent assays: (A) firefly luciferase and (B) mCherry fluorescence. The dose rate for all VCLA irradiations was 0.17 Gy/s. As supplement to Figure 4, here both the terminal luminescence and final-day fluorescence data was reordered to demonstrate direct tumor control comparison on the VCLA vs. the HIGS for each paired dose. Significance was determined using either a standard t-test (for comparisons between two groups) or a One-way ANOVA using a Brown-Forsythe and Welch correction (for comparisons between three or more groups).

Cytokine expression fold change of P9 350 µm hemi-coronal Sprague Dawley rat brain slices three days post-irradiation with either a Varian clinical linear accelerator (VCLA) or the novel HIGS-FLASH beam. On Day 3 post-RT, each brain slice was flooded with 250 µL 1X phosphate buffered saline (PBS) without calcium or magnesium at 37C with 5% CO2 in a tissue culture incubator. 100 µL of this collected PBS “rinse” was then sent to Eve Technologies in Calgary, Canada for assessment using “RD27” Millipore Sigma cat #RECYTMAG-65K to measure the expressed levels of 27 different selected cytokines. A PBS-only blank control was included. To make the heat map demonstrating cytokine signal fold change for each dose rate, raw values were averaged and normalized against their paired 0 Gy control after the (negligible) value for the PBS-only blank was subtracted. These fold changes were then assembled in Prism GraphPad for a representation of which cytokines expressed more or less with dose rates of 0.17 Gy/s (conventional VCLA), 0.79 Gy/s (HIGS sub-FLASH), and 54.02 Gy/s (HIGS FLASH).

**SFigure 5:**
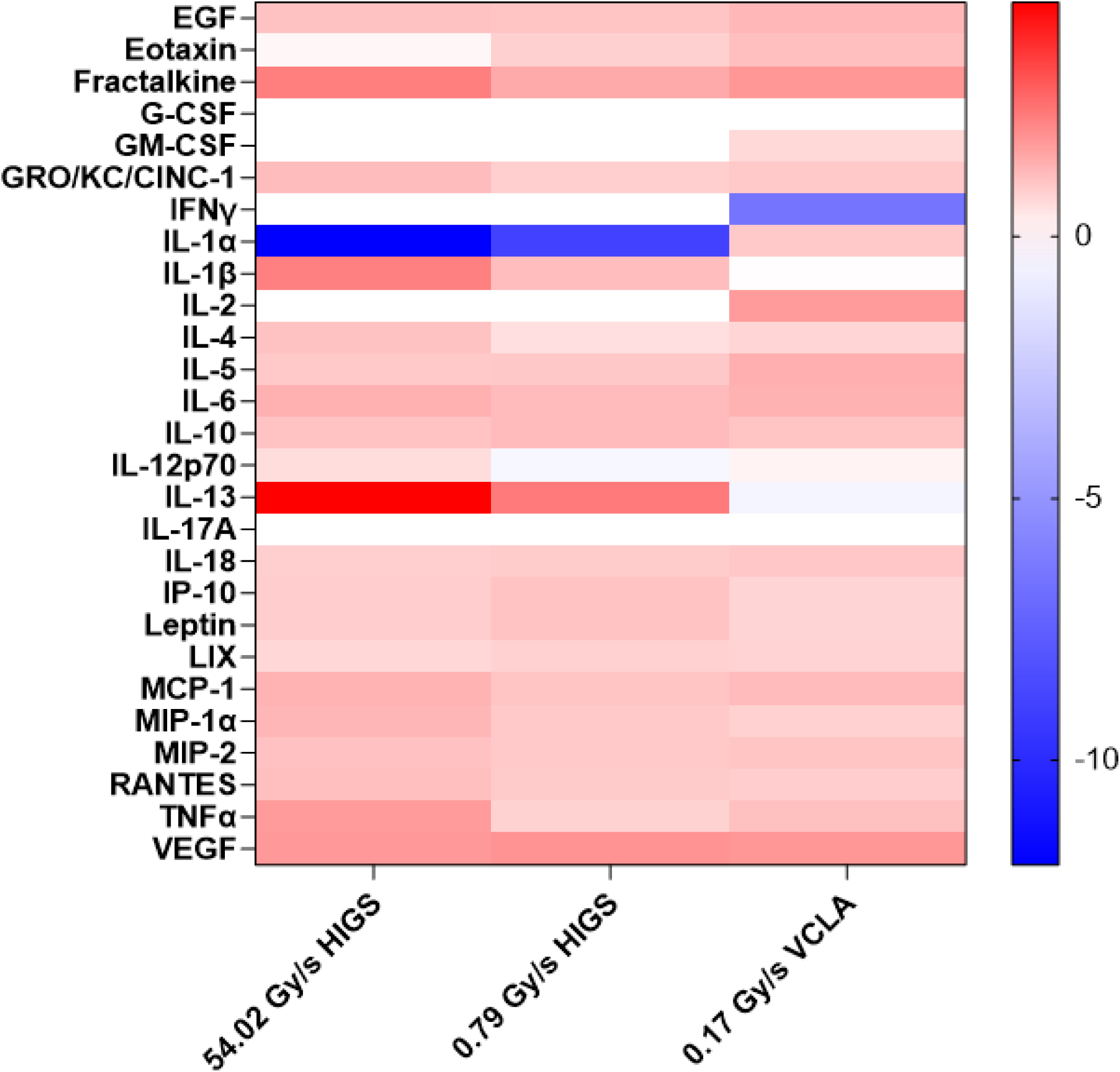
Heat map demonstrating fold-change of cytokine expression of rat brain slices exposed to ∼24.31 Gy at dose rates of 0.17, 0.79, and 54.02 Gy/s using either the VCLA or the HIGS-FLASH beam, respectively. PBS rinses from n = 4 - 8 brain slices were profiled for 27 different rat cytokines. Raw values per MDR were averaged and normalized against their associated 0 Gy control as a fold change. Specifically, the FLASH dose rate of 54.02 Gy/s demonstrated significant increase of Fractalkine, IL-1β, IL-13, and TNFα over both sub-FLASH dose rate of 0.79 Gy/s and conventional dose rate of 0.17 Gy/s. Increase in Fractalkine specifically is linked to increase in microglial activation, of which can be seen in Fig6.

## Notes

### Competing Interest Statement

Zachary J. Reitman has received royalties for intellectual property related to brain tumor diagnostic tests that is managed by Duke and has been licensed to Genetron Health, and honoraria from Eisai Pharmaceuticals and Oakstone Publishing. Scott R. Floyd is on the board of directors and has stock in Round Table Research, Inc.

